# Triacylglycerol fatty acid composition modulates ATGL regioselectivity and diacylglycerol output

**DOI:** 10.64898/2026.10.09.757750

**Authors:** Nadine B. Egger, Thomas Züllig, Christian Gruber, Karl Gruber, Franz P. W. Radner, Amit Singh Sahrawat

**Affiliations:** Institute of Molecular Biosciences, University of Graz, Graz, Austria; BioTechMed-Graz, Graz, Austria; Field of Excellence BioHealth, University of Graz, Graz, Austria

## Abstract

Adipose triglyceride lipase (ATGL) catalyses the first and rate-limiting step of intracellular triacylglycerol (TAG) hydrolysis, yielding diacylglycerol (DAG) and a free fatty acid (FA). The site of TAG cleavage determines the regioisomeric identity of the DAG product, with distinct consequences for downstream signalling and lipid metabolism. Here, we show that ATGL prefers specific fatty acyl chain lengths and that this preference contributes to its regioselectivity on heterogeneous TAG substrates. Structural modelling identified a substrate-binding pocket comprising a hydrophilic anchor and a hydrophobic tunnel whose dimensions favour long-chain FAs. Mass spectrometry-based analysis of DAG products generated by ATGL from defined TAG substrates revealed an intrinsic preference for terminal (*sn*-1/*sn*-3) ester bond cleavage. Acyl-chain compatibility modulated this preference, with poorly accommodated terminal chains increasing the relative contribution of *sn*-2 cleavage. Consequently, ATGL generated predominantly *sn*-2-retaining DAG species from heterogeneous TAG substrates containing long-chain fatty acyl chains, including substantial amounts of the *sn*-1,2-DAG regioisomer, which accounted for 82% of the DAG products generated from the defined TAG species 18:0/18:1/16:0. These findings establish TAG fatty acyl composition as a determinant of ATGL regioselectivity, thereby modulating the regioisomeric profile of ATGL-derived DAGs and their potential contribution to downstream lipid metabolism, including phospholipid synthesis.

## Introduction

Triacylglycerols (TAGs) constitute the major energy reserve in mammals, storing excess fatty acids (FAs) in a metabolically inert form within intracellular lipid droplets.^1–4^ Each TAG molecule consists of three fatty acyl chains esterified at specific positions on a glycerol backbone (*sn*-1, *sn*-2, and *sn*-3). As a result, TAGs comprise a chemically diverse lipid pool with fatty acyl chain composition and positional distribution varying widely.^5–8^ While lipid droplets primarily contain TAG species rich in long-chain fatty acids (FAs, C13-C21), their TAG composition can also include a diverse range of short (<C6), medium- (C6-C12), and very-long-chain (≥C22) FAs.^9–16^

Upon increased metabolic demand, TAGs undergo sequential hydrolysis, collectively termed lipolysis, to release FAs for beta-oxidation or other metabolic fates.^17^ The first and rate-limiting step in this process is catalysed by adipose triglyceride lipase (ATGL, also known as patatin-like phospholipase domain-containing protein 2, PNPLA2; UniProt Q96AD5), which hydrolyses a single ester bond within the TAG molecule to release one free FA and generate diacylglycerol (DAG).^17–19^ In addition to its role as an intermediate in TAG breakdown, DAG has important functions in cellular lipid metabolism. In this context, a recent study by Wijesinghe et al. demonstrated that ATGL activation remodels cellular DAG pools and promotes phospholipid synthesis, highlighting a link between ATGL activity, DAG metabolism, and membrane lipid synthesis.^20^

Efficient TAG hydrolysis requires the ATGL co-activator alpha/beta hydrolase domain-containing protein 5 (ABHD5, also known as comparative gene identification-58, CGI-58).^17,21^ The physiological importance of this pathway is illustrated by genetic studies: ATGL deficiency in mice causes massive TAG accumulation, cardiac lipotoxicity, and premature death,^22^ whereas loss-of-function variants in humans result in neutral lipid storage disease with myopathy.^23^ Conversely, excessive ATGL activity promotes pathological release of free FAs and contributes to insulin resistance, hepatic steatosis, cardiovascular disease, and neuroinflammation.^17,24,25^ These observations establish ATGL as a central regulator of lipid homeostasis and an attractive therapeutic target.^18,26,27^

The metabolic and signalling consequences of ATGL-mediated lipolysis may therefore depend not only on the amount of DAG generated but also on its molecular identity. For DAG, this identity is determined by the site of TAG ester bond cleavage: hydrolysis at *sn*-1, *sn*-2, or *sn*-3 yields *sn*-2,3-, *sn*-1,3-, or *sn*-1,2-DAG, respectively (Fig. 1a). Among these regioisomers, *sn*-1,2-DAG is the canonical endogenous ligand for protein kinase C (PKC)^28,29^ and the immediate precursor for phospholipid synthesis via the CDP-choline/ethanolamine (Kennedy) pathway^30,31^. In contrast, *sn*-1,3-DAG and *sn*-2,3-DAG are primarily subject to further metabolic processing.^32,33^ Thus, the DAG regioisomer generated by ATGL links the site of TAG hydrolysis to distinct downstream metabolic and signalling pathways.^22,24,25,34–37^

**Figure 1.**
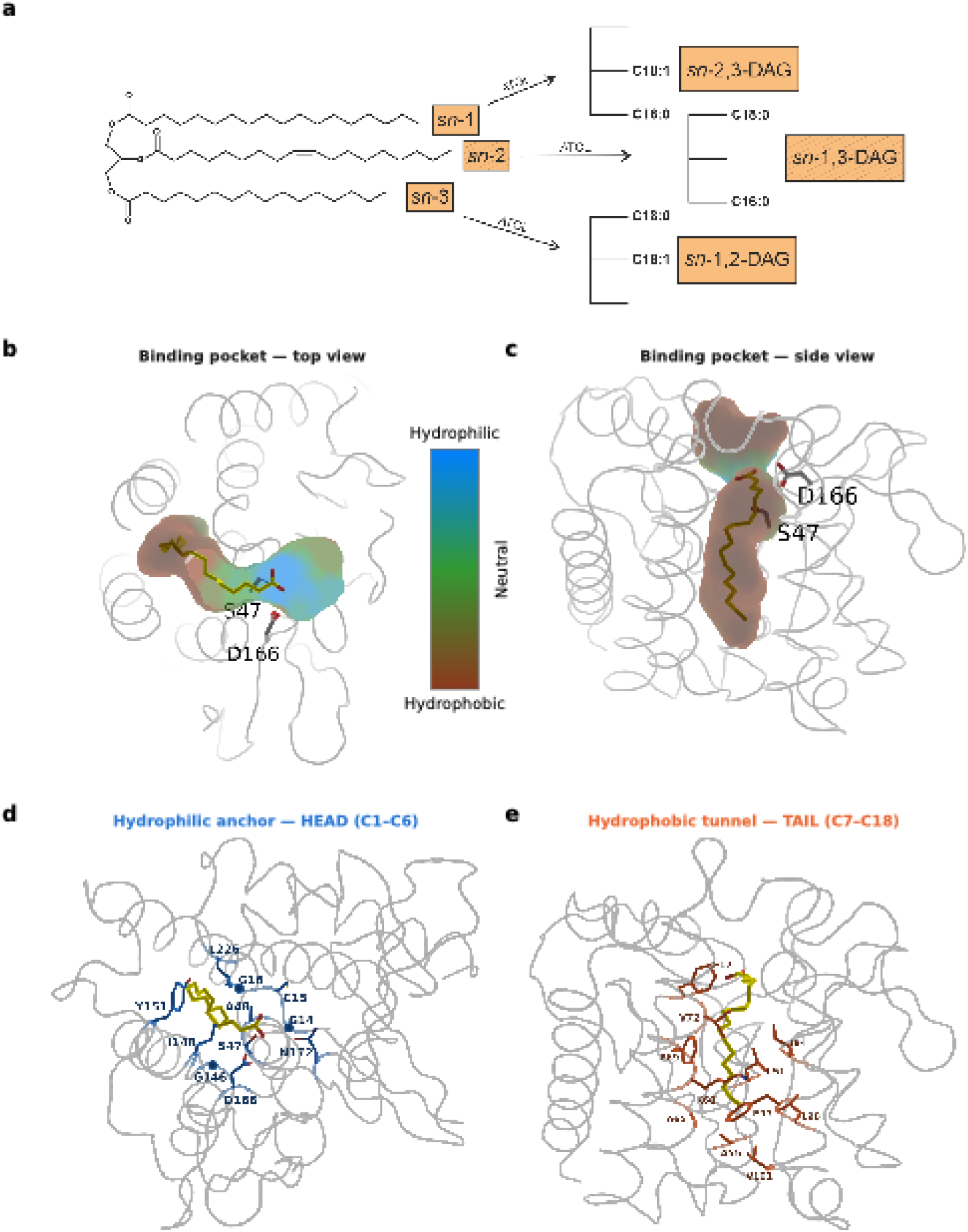
The ATGL substrate-binding pocket comprises a hydrophilic anchor and a hydrophobic acyl tunnel. (**a**) Structural basis of ATGL regioselectivity: Cleavage of a TAG molecule at *sn*-1, *sn*-2, or *sn*-3 position generates *sn*-2,3-DAG, *sn*-1,3-DAG, or *sn*-1,2-DAG, respectively. The resulting DAG regioisomers differ in their metabolic and signalling properties: *sn*-1,2-DAG activates protein kinase C and serves as substrate for the Kennedy-pathway, whereas *sn*-1,3-DAG and *sn*-2,3-DAG have distinct metabolic fates. (**b**, **c**) Structure of the ATGL substrate-binding pocket with FA 18:0 shown as yellow sticks and the catalytic residues S47 and D166 indicated. The cavity surface is coloured according to the CavitOmiX hydrophobicity potential (shared scale: blue, hydrophilic; brown, hydrophobic), viewed from the top (**b**) and the side (**c**). The pocket comprises a hydrophilic anchor near the catalytic centre connected to a predominantly hydrophobic acyl tunnel. (**d**) The hydrophilic anchor accommodates the carboxyl-proximal region (C1–C6) of the FA. Residues identified as FA 18:0 contact sites in MD simulations are shown as ribbons: G14, C15, G16, S47, A48, G146, I148, D166, N172, L226, and Y151. Contact was defined as heavy-atom distances ≤4.0 Å with an occupancy ≥30%. (**e**) The hydrophobic tunnel accommodates the distal region of the FA acyl chain (C7–C18). Contact residues include F17, L51, A55, A65, K68, F69, V72, I94, F97, L98, and V101, with F69 showing the highest contact occupancy. The geometry of this tunnel is predicted to contribute to fatty acyl chain-length recognition.

Despite the emerging role of ATGL-derived DAG in cellular lipid metabolism, the molecular basis of ATGL substrate recognition and regioselectivity remains poorly understood. In particular, it remains unclear how ATGL selects which of the three ester bonds in a TAG molecule to hydrolyse, thereby determining the regioisomeric identity of the resulting DAG and its potential metabolic fate. No experimental structure of ATGL is currently available, and the architecture of its substrate-binding pocket has not been described. To date, systematic biochemical analysis of ATGL regioselectivity has been limited to a single substrate, triolein (18:1/18:1/18:1), in which the *sn*-2 position was identified as the preferred site of hydrolysis, with broader activity towards *sn*-1 upon co-activation by ABHD5.^33^ No turnover was detected at *sn-*3, leading to the widely accepted view that ATGL primarily generates *sn*-1,3-DAG. Triolein, however, does not reflect the combinatorial heterogeneity of the physiological lipid droplet TAG pool,^5,7,8^ in which chemically distinct fatty acyl chains occur in different combinations and occupy different *sn*-positions of the glycerol backbone. A substrate-binding pocket that prefers particular fatty acyl chain lengths could therefore influence not only substrate recognition but also the *sn*-position of ester bond cleavage. Whether such a link between chain-length preference and regioselectivity shapes the DAG regioisomer profile generated from physiologically relevant TAG substrates remains unknown.

Here, we demonstrate that fatty acyl chain length is a key determinant of ATGL regioselectivity on heterogeneous TAG substrates. Using AI-assisted structural modelling, we characterise the architecture of the ATGL substrate-binding pocket, revealing a conserved hydrophilic anchor and a hydrophobic acyl tunnel whose dimensions favour long-chain FAs. We show that fatty acyl chain composition modulates ATGL cleavage-site preference in heterogeneous TAG substrates, favouring hydrolysis of the *sn*-1 and *sn*-3 ester bonds when the corresponding acyl chains are compatible with the substrate-binding pocket. Quantitative ultra-high-performance liquid chromatography-tandem mass spectrometry (UHPLC-MS/MS) analysis of DAG regioisomer formation further demonstrates that this substrate-dependent shift in cleavage-site preference redirects ATGL product output towards *sn*-2-retaining DAG species, including the signalling-competent *sn*-1,2-DAG regioisomer. Together, these findings establish ATGL as a fatty acyl chain length-dependent regioselective lipase and provide a mechanistic link between TAG molecular diversity and the DAG regioisomer output of lipolysis, with implications for the generation of DAG species involved in phospholipid synthesis and lipid signalling.

## Results

### ATGL contains an amphiphilic substrate-binding pocket

To determine whether ATGL contains a defined substrate-binding pocket capable of discriminating fatty acyl chain lengths, we generated substrate-bound structural models of the catalytic patatin domain of ATGL (residues 10-179; PROSITE: PRU01161) in complex with saturated FAs spanning the physiological range of fatty acyl chain lengths found in intracellular TAG pools (C4:0-C28:0). We used free FAs as a simplified system to isolate chain-length complementarity from the additional structural constraints arising from the glycerol backbone and the other two fatty acyl chains. Across the FA series, the models consistently positioned the carboxylate group adjacent to the catalytic S47/D166 dyad within a single binding cavity, while the acyl moiety extended into the cavity in an orientation compatible with catalysis. The cavity thus comprises two functionally distinct regions: a hydrophilic anchor that engages the FA carboxylate group and a hydrophobic tunnel that accommodates the fatty acyl chain (Fig. 1b and c).

The hydrophilic anchor positions the FA carboxylate through interactions with residues surrounding the catalytic site (Fig. 1d), while additional residues in this region contact the proximal portion of the fatty acyl chain (C1-C6), further stabilising substrate orientation (Supplementary Fig. S1a). Extending from this anchor, a larger hydrophobic tunnel accommodates the remaining fatty acyl chain (Fig. 1e). This tunnel is lined predominantly by aromatic and hydrophobic residues, which together form a complementary surface for the aliphatic hydrocarbon chain (Supplementary Fig. S1b). Together, these features define an amphiphilic substrate-binding pocket in which the polar carboxylate group is positioned near the catalytic site while the hydrophobic acyl chain extends into a non-polar tunnel.

The pocket architecture is highly conserved across ATGL orthologues,^38^ and several of the identified residues have previously been implicated in substrate or inhibitor binding by mutagenesis.^38,39^ Notably, co-folded structural ensembles recovered a substrate-binding pocket without prior specification, providing independent support for the modelled architecture.^40–42^ We next asked how fatty acyl chain length affects the complementarity of the acyl tunnel.

### The geometry of the ATGL substrate-binding pocket favours long-chain fatty acids

We first used all-atom molecular dynamics (MD) simulations to determine how FAs of varying chain lengths occupy the hydrophobic tunnel. Anchoring the carboxylate group near the catalytic residue S47 allowed us to compare the conformational behaviour and spatial occupancy of FA chains across the binding pocket.

The tunnel dimensions favoured long-chain FAs. Short- and medium-chain FAs (C4:0-C12:0) did not fully occupy the distal portion of the tunnel, whereas long-chain FAs established maximal contact with the buried hydrophobic residues forming the pocket floor (Fig. 2a and b). The tunnel is not linear but contains a distinct bend at the transition between the hydrophilic anchor and the hydrophobic tunnel. Long-chain FAs accommodated this geometry by adopting defined conformations within the binding pocket, whereas very-long-chain FAs (≥C22:0) exceeded the available tunnel length and required additional bending to fit within the cavity (Fig. 2c and d). Quantitative analysis of ligand elongation and pocket-packing efficiency supported these observations (Supplementary Fig. S2). Ligand elongation plateaued beyond the long-chain range, indicating that additional carbon atoms in very-long-chain FAs were accommodated through chain bending rather than further extension along the hydrophobic tunnel. Similarly, pocket-packing efficiency increased from 64% for short-chain FAs to 85% for long-chain FAs and did not increase further for longer chains. Thus, long-chain FAs achieved the highest degree of occupancy of the available hydrophobic tunnel space, whereas shorter and very-long-chain FAs showed reduced geometric complementarity.

**Figure 2.**
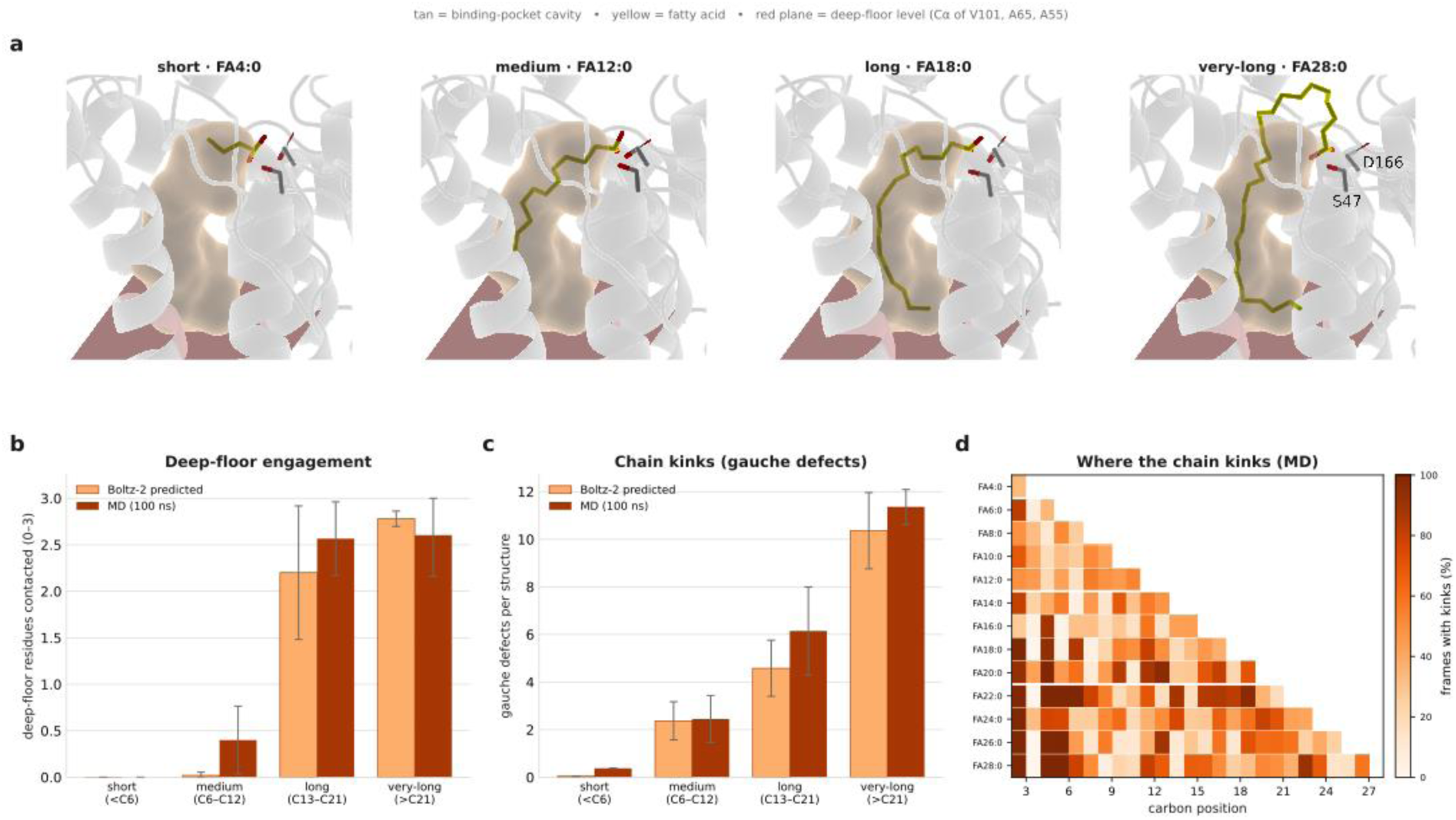
Long-chain fatty acids achieve maximal occupancy of the ATGL acyl tunnel. (**a**) Representative Boltz-2 predicted ATGL-FA complexes for FA 4:0, FA 12:0, FA 18:0, and FA 28:0. The substrate-binding cavity is shown in tan, FAs are shown as sticks (yellow carbon atoms, red oxygen atoms), and the catalytic residues S47 and D166 are shown in grey. The semi-transparent red plane marks the "deep floor" of the tunnel, defined by a plane through the Cα atoms of V101, A65, and A55. The acyl chain remains above the pocket floor for FA 4:0), reaches the floor for FA 18:0, and bends back within the cavity for FA 28:0.(**b**) Deep-floor engagement, defined as the number of the three deep-floor residues (A55, A65, and V101) with at least one heavy atom within 4.5 Å of any ligand carbon. Light orange denotes Boltz-2-predicted ensembles (25 models per FA), and dark orange denotes MD trajectories. Deep-floor engagement increased with acyl chain length and plateaued in the long-chain range. (**c**) Number of acyl-chain kinks, defined as gauche defects according to the indicated torsional-angle criterion (|τ| ≤120°), across MD trajectories. (**d**) Distribution of gauche defects along the acyl chain. Cell colour indicates the percentage of MD frames in which the corresponding C-C bond adopted a gauche confirmation.

We next asked whether this geometric preference was also reflected energetically. Using a free-energy perturbation (FEP) protocol, we stepwise extended the FA alkyl chain from C4:0 to C24:0 and calculated the relative binding free energy for each additional ethyl group. Tail extension remained energetically favourable from short- to long-chain FAs but became unfavourable beyond this range (Supplementary Fig. S3). Thus, the energetic profile provided independent support for the optimum identified from the MD ensembles and spatial analyses.

As an independent and higher-throughput measure of chain-length compatibility, we next evaluated the binding-pose metrics generated by the co-modelling approach, which provide a score for the predicted compatibility of each FA binding pose within the ATGL substrate-binding pocket, with lower *F*_score_ values indicating better compatibility. The resulting *F*_score_ values decreased progressively from C4:0 to C16:0, reached a broad minimum across C16:0-C18:0, and increased again for longer FAs (Supplementary Fig. S4). This analysis, therefore, identified C16:0-C18:0 as the range with the highest predicted compatibility with the ATGL substrate-binding pocket, consistent with the MD, packing, and FEP analyses.

Together, these complementary analyses show that the geometry of the hydrophobic tunnel favours long-chain FAs, whereas shorter and very long-chain FAs exhibit less favourable geometric complementarity. Short-chain FAs fail to establish sufficient hydrophobic interactions within the substrate-binding pocket, whereas very-long-chain FAs exceed its optimal dimensions. This structural organisation of the ATGL substrate-binding pocket provides a plausible mechanism by which the fatty acyl chain composition of a heterogeneous TAG molecule may influence which ester bond is positioned for hydrolysis. Because the hydrophobic tunnel accommodates a single fatty acyl chain (Fig. 1b and c), the compatibility of each fatty acyl chain with the tunnel could influence which of the three ester bonds is positioned favourably for attack by the catalytic S47. In a heterogeneous TAG, this would be expected to favour hydrolysis of the ester bond associated with the most compatible fatty acyl chain. Accordingly, TAG species containing compatible long-chain fatty acyl chains at the terminal *sn*-1 or *sn*-3 positions, together with a less compatible short-or very-long-chain fatty acyl chain at *sn*-2, are predicted to favour terminal ester bond cleavage.

### ATGL preferentially hydrolyses the *sn*-1 and *sn*-3 ester bonds in heterogeneous TAG substrates

To directly test whether fatty acyl chain composition determines ATGL regioselectivity, we quantified DAG regioisomer formation from chemically defined TAG substrates by UHPLC-MS/MS. Because each substrate contained defined FAs at defined *sn*-positions, the acyl composition of the resulting DAG products directly indicated which ester bond was hydrolysed by ATGL. TAG hydrolysis assays were performed using purified human ATGL (Supplementary Fig. S5) either alone or in the presence of the co-activator ABHD5.

The fully asymmetric, stereochemically defined TAG 18:0/18:1/16:0 served as an informative first substrate because cleavage at each *sn-*position generates a unique DAG product, allowing direct determination of the hydrolysis site from the DAG acyl composition without requiring chiral separation (Fig. 3a). Using this substrate, ATGL preferentially cleaved the *sn*-1 and *sn*-3 ester bonds, which together accounted for 99% of total DAG formation, whereas cleavage at the *sn*-2 position contributed only 1%. The *sn*-1 and *sn*-3 ester bonds were hydrolysed at a ratio of approximately 1:5, indicating a strong preference for *sn*-3 over *sn*-1 cleavage and, overall, a pronounced discrimination against the central *sn*-2 position. Consequently, 82% of the total DAG generated corresponded to the signalling-competent *sn*-1,2-DAG regioisomer, which was not detected during triolein hydrolysis under similar conditions.^33^ Co-activation by ABHD5 increased total TAG hydrolysis, as reflected by higher DAG formation, and caused a modest shift towards *sn*-2 cleavage, while terminal *sn*-1/*sn*-3 ester bonds remained the predominant sites of hydrolysis.

**Figure 3.**
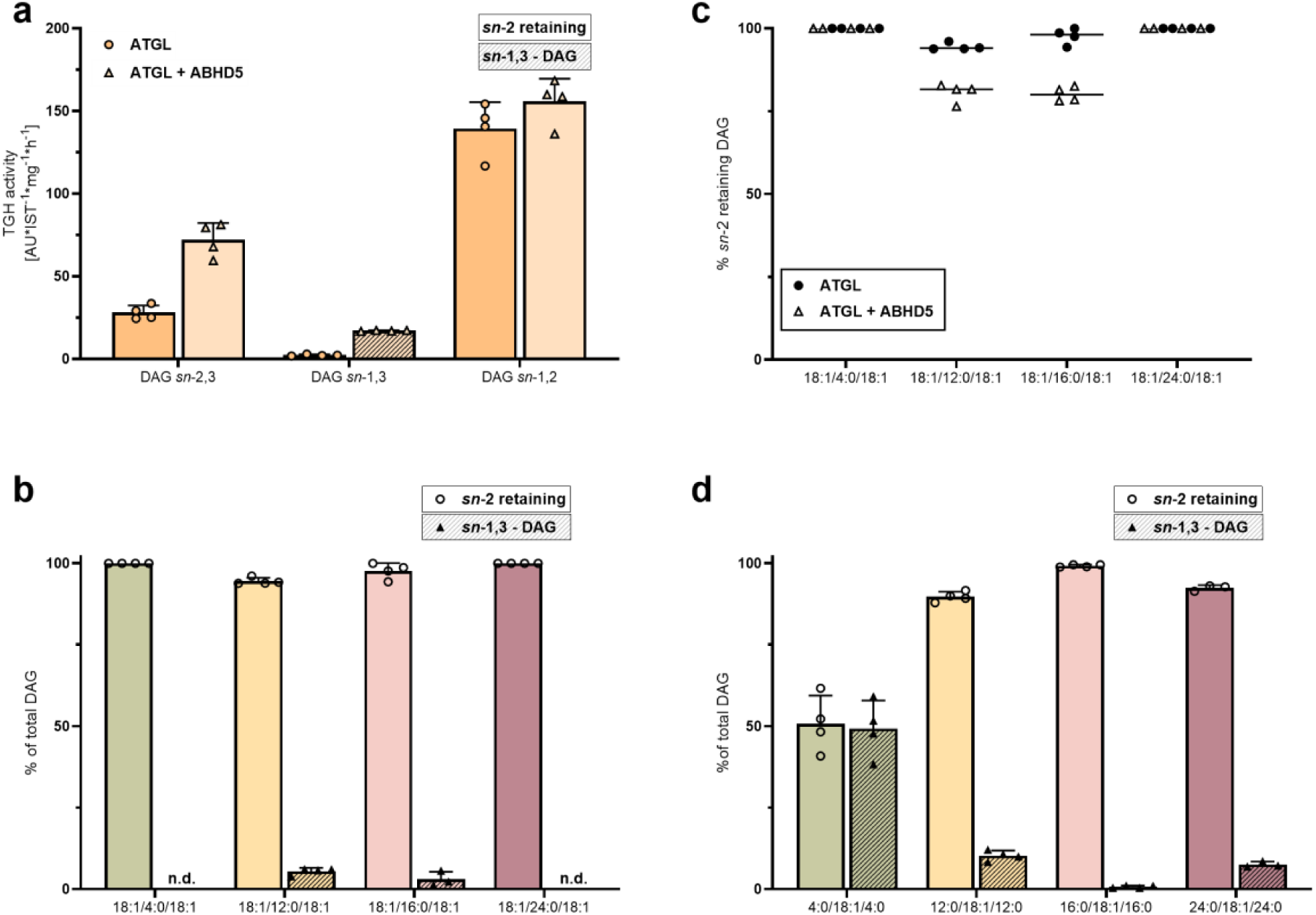
Fatty acyl chain composition determines ATGL regioselectivity on heterogeneous TAG substrates. DAG products generated from defined TAG substrates were quantified by UHPLC-MS/MS analysis following incubation with purified human ATGL in the absence or presence of ABHD5. DAG products are grouped according to the site of ester bond cleavage as *sn*-2-retaining DAG (generated by *sn*-1/*sn*-3 cleavage) or *sn*-1,3-DAG (generated by *sn*-2 cleavage). In panels a, b and d, hatched bars denote *sn*-1,3-DAG, whereas open bars denote *sn*-2-retaining DAG species. Individual data points are shown as circles and triangles, with their meaning indicated in the corresponding panels. Data are presented as mean ± SD (n = 4). (**a**) DAG regioisomer formation from the asymmetric, stereochemically defined TAG substrate 18:0/18:1/16:0. The unique fatty acyl composition enables direct determination of the hydrolysed ester bond without chiral separation. TAG hydrolase (TGH) activity is expressed as arbitrary units normalized to the internal standard per mg protein per hour (AU IST^-1^ mg^-1^ protein h^-1^). Light and dark bars indicate ATGL alone and ATGL + ABHD5, respectively. (**b**) Relative abundance of *sn*-2-retaining DAG and *sn*-1,3-DAG generated by ATGL from the symmetric TAG series 18:1/X/18:1 (X = 4:0, 12:0, 16:0, or 24:0), expressed as percentage of total DAG. n.d., not detected. (**c**) Comparison of *sn*-2-retaining DAG formation from the symmetric TAG series 18:1/X/18:1 (X = 4:0, 12:0, 16:0, or 24:0) in the presence or absence of ABHD5. The percentage of *sn*-2-retaining DAG relative to total DAG is shown. Horizontal lines indicate mean values, and individual measurements are shown as data points. (**d**) Relative abundance of *sn*-2-retaining DAG and *sn*-1,3-DAG generated by ATGL from the reciprocal TAG series X/18:1/X (X = 4:0, 12:0, 16:0, or 24:0), expressed as percentage of total DAG.

We next examined whether TAG fatty acyl chain composition and *sn*-position-specific fatty acyl chain distribution influence ATGL regioselectivity. In a symmetric TAG series (18:1/X/18:1; X = 4:0, 12:0, 16:0, or 24:0), in which the flanking positions were esterified with oleate and the central acyl chain was varied, ATGL consistently generated predominantly *sn*-2-retaining DAG species. The corresponding *sn*-1,3-DAG product resulting from *sn*-2 cleavage remained minor or undetectable. Across all substrates, *sn*-2-retaining DAG accounted for 95-100% of the total DAG generated by ATGL (Fig. 3b). Co-activation by ABHD5 caused a modest increase in *sn*-2 cleavage for C12:0- and C16:0-containing substrates but did not alter the predominant formation of *sn*-2-retaining DAG across the series (Fig. 3c).

To distinguish whether regioselectivity was driven by fatty acyl chain positioning or by an intrinsic preference of ATGL for terminal ester bond cleavage, we analysed a reciprocal TAG series in which oleate was esterified at the *sn*-2 position (X/18:1/X, X = 4:0, 12:0, 16:0, or 24:0). In these substrates, the central long-chain fatty acyl chain was predicted to exhibit the highest complementarity to the hydrophobic tunnel. However, ATGL still preferentially cleaved the terminal *sn*-1 and *sn*-3 ester bonds whenever the flanking fatty acyl chains were compatible with the substrate-binding pocket (Fig. 3d). This preference was most pronounced for C16:0 in this series, as increasing the flanking fatty acyl chains from C12:0 to C16:0 resulted in essentially exclusive *sn*-1/*sn*-3 ester bond cleavage. *sn*-2 cleavage increased again in the C24:0-flanked substrate, in which the flanking fatty acyl chains exceeded the optimal dimensions of the hydrophobic tunnel. Conversely, shortening the flanking fatty acyl chains to C4:0 resulted in an approximately equal distribution between terminal and *sn*-2 ester bond cleavage, as the C4:0 chains were insufficient to efficiently engage the substrate-binding pocket. These findings demonstrate that ATGL retains an intrinsic preference for terminal ester bond cleavage, while fatty acyl chain complementarity modulates this preference when the flanking fatty acyl chains are poorly accommodated within the substrate-binding pocket.

Together, these experiments establish that ATGL preferentially hydrolyses terminal ester bonds in heterogeneous TAG substrates, thereby predominantly generating *sn*-2-retaining DAG species. This substrate-dependent regioselectivity contrasts with the classical triolein paradigm and identifies fatty acyl chain composition as a key determinant of ATGL product profile.

## Discussion

Our findings reveal that ATGL substrate recognition is shaped by a defined substrate-binding pocket that favours specific fatty acyl chain lengths. Structural modelling identified a hydrophilic anchor and a hydrophobictunnel, whose dimensions best match long-chain FAs. This architecture provides a structural explanation for the previously reported long-chain preference of ATGL.^18,33^ It also illustrates why triolein has been a useful model substrate: its three 18:1 acyl chains fall within the optimal range of the ATGL binding pocket. More broadly, our findings indicate that ATGL does not recognise TAGs solely as defined molecular species but can accommodate individual fatty acyl chains within a preferred chain-length range. One possible interpretation is that such recognition provides an efficient solution to the combinatorial diversity of mammalian TAGs, which comprise numerous molecular species that differ in their acyl chain composition and positional distribution.^5,7–9^ Recognition of the individual acyl chain at the reacting position, rather than the entire TAG molecular species, may therefore allow ATGL to accommodate a diverse physiological substrate pool while retaining a preference for specific chain lengths. Importantly, this mode of substrate recognition also provides a plausible mechanism for linking fatty acyl chain compatibility to the selection of the ester bond that is hydrolysed, thereby connecting TAG composition with the DAG regioisomer output of lipolysis.

Importantly, our findings extend the understanding of ATGL regioselectivity beyond the traditional triolein model.^33^ Triolein carries three identical FA 18:1 chains and therefore does not reflect the structural heterogeneity of the mammalian TAG lipidome, in which dominant species carry distinct long-chain fatty acyl chains and the *sn*-2 position is typically more unsaturated than the terminal positions.^5,9^ The fully asymmetric, stereochemically defined TAG 18:0/18:1/16:0 captures these features more closely. With this substrate, ATGL preferentially hydrolysed the terminal ester bonds, whereas cleavage at *sn*-2 was minor. This intrinsic preference for the terminal ester bonds was evident across heterogeneous TAG substrates whenever the terminal fatty acyl chains were compatible with the substrate-binding pocket. The reciprocal TAG series further showed that regioselectivity was not simply determined by the presence of the best-fitting long-chain acyl group placed at *sn*-2. Even when FA 18:1 occupied the *sn*-2 position and was predicted to provide the highest complementarity to the hydrophobic tunnel, ATGL retained its preference for terminal ester bond cleavage as long as the flanking fatty acyl chains were compatible with the pocket. Fatty acyl chain compatibility modulated this intrinsic preference: poor accommodation of the terminal chains increased the relative contribution of *sn*-2 cleavage, whereas compatible terminal fatty acyl chains supported predominantly *sn*-1/*sn*-3 hydrolysis. Thus, ATGL regioselectivity reflects the interplay between an intrinsic preference for terminal ester bond cleavage and the compatibility of the acyl chains occupying the substrate-binding pocket. A similar preference for hydrolysis of terminal ester bonds has been reported for PNPLA3, which shares the catalytic patatin domain with ATGL.^43^ These findings suggest that substrate composition should be considered when interpreting regioselectivity across the PNPLA enzyme family and, more broadly, in studies of lipase substrate specificity.

The site of ATGL cleavage has consequences beyond the identity of the hydrolysed ester bond, as it determines the regioisomeric composition of the resulting DAG pool. In the fully asymmetric, stereochemically defined TAG, preferential *sn*-3 cleavage resulted in predominantly *sn*-1,2-DAG, demonstrating that ATGL can generate substantial amounts of this signalling-competent and phospholipid-precursor DAG regioisomer. Because *sn*-1,2-DAG is a canonical endogenous ligand for PKC^28,29^ and has been specifically implicated in lipid-induced insulin resistance,^24,25,44–46^ our findings raise the possibility that substrate-dependent ATGL regioselectivity contributes to the molecular composition of cellular DAG pools.

The potential metabolic relevance of this finding is supported by a recent study by Wijesinghe et al., which demonstrated that ATGL activation remodels cellular DAG pools and promotes phospholipid synthesis.^20^ That study highlighted the contribution of ATGL-mediated lipolysis to membrane lipid metabolism, but the molecular basis for generating distinct DAG species remained unresolved. Our findings provide a mechanistic explanation by showing that the TAG fatty acyl composition modulates the regioisomeric identity of ATGL-generated DAGs. Thus, substrate-dependent ATGL regioselectivity may shape the composition of DAG pools available for downstream metabolic pathways, including phospholipid synthesis.

This finding may also be relevant to the recycling of acylglycerols into phospholipids. Igal and Coleman reported impaired acylglycerol recycling from TAG to phospholipids in fibroblasts from patients with neutral lipid storage disease associated with loss of functional ABHD5, despite preserved lipase activity.^47,48^ This observation suggested that defective acylglycerol recycling, rather than impaired TAG hydrolysis, contributes to altered phospholipid metabolism in these cells. Our findings raise the possibility that variation in the regioisomeric composition of ATGL-derived DAG could influence the pool of DAG available for phospholipid synthesis, providing a potential link between TAG substrate composition, ATGL regioselectivity, and phospholipid metabolism. Whether altered ATGL regioselectivity contributes to the phospholipid abnormalities associated with neutral lipid storage disease remains to be determined.

The second product of ATGL-mediated hydrolysis is a free FA, and FA chain length can likewise influence cellular responses, including insulin resistance, inflammation, apoptosis, and immunity.^37,49–55^ Thus, TAG molecular composition has the potential to influence the molecular identity of both products generated in the first step of lipolysis. This consideration has implications for quantitative models of lipid metabolism, which commonly represent lipolysis as a sequential conversion between lumped TAG, DAG, and MAG pools or are parameterised using triolein-like substrates.^56–58^ Such representations do not account for substrate-dependent differences in DAG regioisomer or FA output. The chain-length compatibility score and experimentally determined regioselectivity ratios generated here could provide relative parameters for incorporating substrate-dependent ATGL activity, including differences in DAG regioisomer and FA output, into more species-resolved models of lipolysis and its downstream metabolic and signalling pathways.

From a structural and kinetic perspective, several mechanisms could account for the intrinsic regioselectivity of ATGL. First, the glycerol backbone may adopt conformations in which the *sn*-2 ester is less favourably positioned for attack than the terminal ester bonds. Second, the ATGL binding site may preferentially accommodate TAG conformations that position a terminal ester bond near the active site. Third, differences in the activation energy for ester bond cleavage could contribute to the observed preference, with a higher barrier for *sn*-2 cleavage resulting in a lower hydrolysis rate than for terminal ester cleavage. Distinguishing between these possibilities will require dedicated structural and kinetic analyses. The structural elements of the ATGL binding site identified here may also provide a basis for developing ATGL inhibitors with broader cross-species activity. Current ATGL inhibitors, including atglistatin^26^ and NG-497^39^, target hydrophobic regions of the substrate-binding pocket that exhibit species-specific differences, thereby limiting their cross-species efficacy. The conserved hydrophilic anchor identified in our structural models represents an additional, more conserved feature of the substrate-binding pocket that could be explored as an alternative inhibitor-binding site. Targeting conserved elements of the substrate-binding pocket may therefore provide a complementary strategy for developing ATGL inhibitors with broader cross-species activity.

## Methods

### ATGL254 construct and covalent-constraint co-folding

All computations used the human ATGL (UniProt Q96AD5) 254-residue construct (ATGL254), which contains the complete patatin domain and the catalytic S47/D166 dyad but lacks the disordered C-terminal region involved in lipid-droplet-binding.^59^ A multiple sequence alignment of ATGL254 was generated once using Boltz-2 and reused across all co-folding runs. Boltz-2^60^ was used to generate structural ensembles of ATGL254 in complex with each FA ligand applying a single covalent-bond constraint between the catalytic S47-Oγ and the FA carboxyl carbon. This constraint positions the substrate in a near-attack conformation relative to the catalytic nucleophile. For each FA system, 25 diffusion samples were generated, followed by five affinity module samples and five recycling steps, with physical potentials enabled.

### Molecular Dynamics Simulations

#### Unbiased MD

Starting coordinates were taken from the highest-ranked Boltz-2 co-folded models of ATGL254 in complex with each saturated fatty acid (C4:0–C28:0, 13 even-chain species). Protein atoms were described using the Amber ff19SB force field,^61^ and FA ligands were parameterised using GAFF2 with AM1-BCC partial charges. Complexes were solvated in TIP3P water^62^ with a 12 Å buffer and neutralised with NaCl.^63,64^ To maintain the near-attack geometry between the catalytic nucleophile and the FA carboxyl group while leaving the remainder of the ligand unrestrained, a one-sided flat-bottom restraint was applied between S47-Oγ and the carboxyl carbon (C1), with r₀ = 3.0 Å and k = 1.195 kcal mol⁻¹ Å ⁻². The fatty acyl chain and its interaction with the hydrophobic tunnel were otherwise left unbiased. Simulations were performed in OpenMM 7.7 with CUDA.^65^

Following energy minimisation, each system was heated from 10 K to 300 K over 100 ps and equilibrated for 200 ps under NPT conditions with backbone-position restraints. These restraints were released before the production simulation. Production trajectories were then generated for 100 ns at 300 K, with frames saved every 100 ps. The FA 18:0 system was extended to 500 ns. Across all FA systems, approximately 1.7 µs of unbiased MD simulations were generated.

#### Free Energy Perturbation Calculations

Relative binding free energies were calculated using alchemical thermodynamic integration with softcore potentials following AMBER tutorial A39.^66–69^ The perturbation ladder consisted of sequential ethyl-group additions, transforming the FA series from C4:0 to C24:0. Each transformation was performed in both forward and reverse directions. Homologous FAs were represented as dual-topology residues (L0/L1) sharing an identical core. Softcore masks were applied to the hydrogen removed from Cn (scmask1) and to the newly introduced C(n+1)/C(n+2) carbon atoms and their five hydrogens (scmask2). Both end states were net neutral and therefore required no finite-size charge correction. The starting coordinates for L0 FA 4:0 were taken from the final frame of its 100 ns production trajectory.

For each transformation, binding free energies were computed from separate complex and aqueous legs according to ΔΔG_bind = ΔG_complex − ΔG_aqueous. Each leg was sampled over 11 evenly spaced λ windows using gti_add_sc = 25, scalpha = 0.2, scbeta = 50, with MBAR sampling performed every 2 ps. The thermodynamic-integration region was integrated without bond constraints (noshakemask = ’:L0,L1’, ntf = 1), with a timestep of 1 fs; SHAKE was retained for all other bonds.

For each transformation edge and leg, the system underwent energy minimisation, 200 ps of NVT heating from 10 K to 300 K, and 1 ns NPT equilibration, consisting of 500 ps with positional restraints followed by 500 ps without restraints. This preparation was performed once at λ = 0.5. Individual λ windows were subsequently equilibrated for 200 ps at their respective λ values and simulated in two sequential production segments: 2 × 10 ns for the complex leg and 2 × 5 ns for the aqueous leg. The two production segments were treated as independent MBAR trials. As in the unbiased MD simulations, a one-sided flat-bottom restraint between S47-Oγ and C1 was applied to both L0 and L1 states. For each transformation edge, the final estimate was calculated as the antisymmetric average of the forward and reverse transformations, ½(ΔΔG_fwd − ΔΔG_rev). The complete free-energy perturbation campaign comprised approximately 6.6 µs of simulation time.

### Pocket volume, depth, and packing

The binding-pocket geometry was characterised using pyKVFinder^70^ with ligand-guided cavity detection. The following parameters were used: (probe_in_ = 1.4 Å, probe_out_ = 6.0 Å, grid step = 0.4 Å, ligand cutoff = 5.0 Å, and solvent-excluded surface (SES) definition. For each structural model, pocket volume, pocket area, maximum and mean depth, sphericity, elongation, ligand SES volume, and filled and void volumes were recorded. Packing efficiency (Fig. S2) was calculated as 100*filled/(filled+void).

### Chain-length-fitness scoring

To quickly estimate the compatibility of FAs of different chain lengths with the ATGL substrate-binding pocket, we devised an empirical chain-length fitness score (*F*_score_ ) for each FA:

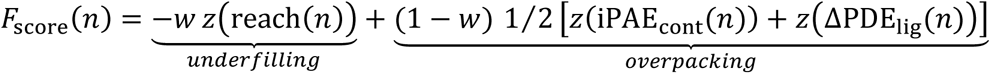

where w=0.5. The first term quantifies underfilling of the substrate-binding tunnel, whereas the second term quantifies overpacking using co-modelling confidence metrics. The fatty acyl chain reach to the tunnel depth was calculated as:

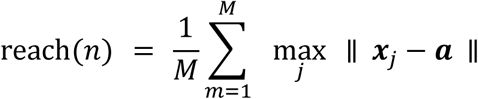

where, for the model *m* of the species *n*, with *x_j_* the ligand heavy-atom positions and *a* the C*α* of the catalytic S47. The reach metrics saturate once the tunnel is occupied. Whereas iPAE_cont_ is the mean PAE over protein–ligand token pairs closer than *R_c_*; *i* runs over protein residue tokens, *j* over ligand atom tokens.

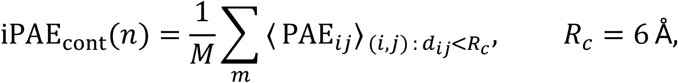

and ΔPDE_lig_ is

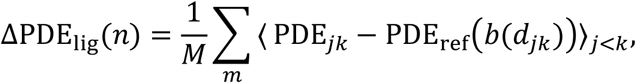

where *j*, *k* are ligand atoms, *b*(*d*) is the 0.5 Å bin containing their separation, and PDE_ref_(*b*) is the mean PDE in that bin pooled over the reference species C4–C12. The short- and medium-chain references have been chosen as they sit in the tunnel with room to spare, so they are conformationally unstrained by construction. Negative values indicate chain geometry better determined than the reference baseline.

Lower *F*_score_ values indicate greater predicted compatibility between the FA and the ATGL substrate-binding pocket.

### Recombinant protein expression and purification

Full-length human ATGL was expressed and purified based on a previously established protocol for PNPLA1 purification (Supplementary Fig. S5),^71^ with modifications adapted from Kulminskaya et al^38^. Expi293F cells (Thermo Fisher Scientific, Waltham, MA, USA) were cultured to a density of 3 × 10⁶ cells ml^-1^ at 37 °C and 125 r.p.m. in a humidified atmosphere containing 7% CO₂, and transfected with 25 µg pcDNA4/HisMaxC-hsATGL^72^ using the ExpiFectamine 293 kit according to the manufacturer’s instructions. Cells were harvested 92 h after transfection, washed three times with ice-cold PBS, and stored at –80 °C until purification.

Cell pellets were resuspended in 50 ml buffer A1 (100 mM potassium phosphate buffer (PPB), pH 7.5, 100 mM KCl, 10% glycerol, 0.1% NP-40, 30 mM imidazole, 1 mM TCEP, 10 mM ATP, 10 mM MgCl_2_, 20 µg ml^-1^ leupeptin, 2 µg ml^-1^ antipain, and 1 µg ml^-1^ pepstatin) and lysed by homogenization (ULTRA-TURRAX®, IKA-Werke, Staufen, Germany; 30 s) followed by sonication (Sonoplus HD 3100, Bandelin, Berlin, Germany; TT13 probe, 25% amplitude, 0.5 s on/off pulses, 3 min) on ice. Lysates were clarified by centrifugation (40,000 × g, 30 min, 4 °C), filtered through a 0.45 µm membrane, and loaded onto a 5 ml HisTrap HP column (Cytiva, Marlborough, MA, USA) connected to an ÄKTA™ pure chromatography system (Cytiva). The column was equilibrated with buffer A1 (7 column volumes (CV)), washed sequentially with buffer A1 (5 CV), high-salt buffer A3 (100 mM PPB, pH 7.5, 500 mM KCl, 10% glycerol, 0.003% NP-40, 30 mM imidazole, and 1 mM TCEP; 7 CV), and buffer A2 (100 mM PPB, pH 7.5, 100 mM KCl, 10% glycerol, and 1 mM TCEP; 10 CV). The column was then washed with 15% buffer B1 (buffer A2 supplemented with 500 mM imidazole; 10 CV), and recombinant ATGL was eluted using a linear gradient to 100% B1 over 5 CV, collecting 2 ml fractions. Fractions were analysed by immunoblotting and Coomassie-stained SDS-PAGE to assess protein identity and purity.

ABHD5 was purified based on previously described protocols.^73,74^ SMT-tagged ABHD5 was co-expressed with the molecular chaperone Tig (encoded by plasmid pTf16; Takara Bio, Kusatsu, Shiga, Japan) in *E*. *coli* BL21(DE3). Cultures were grown at 37 °C in the presence of 50 µg ml^-1^ kanamycin and 34 µg ml^-1^ chloramphenicol, and chaperone expression was induced by addition of 1 g l^-1^ L-arabinose. At an OD_600_ of 0.6, SMT-ABHD5 expression was induced with 0.5 mM IPTG, followed by incubation for 9-12 h at 30 °C.

Cells were lysed by sonication in 50 ml wash buffer (20 mM Tris-HCl, pH 7.8, 500 mM NaCl, 30 mM imidazole, 0.1% NP-40, and 3.5 mM β-mercaptoethanol) supplemented with EDTA-free protease inhibitors (cOmplete, Roche, Basel, Switzerland), 750 U benzonase nuclease (Merck, Darmstadt, Germany), and 1 mg ml^-1^ lysozyme. Following clarification by centrifugation (30,000 × g, 40 min), SMT-ABHD5 was purified using a 5 ml HisTrap HP column (Cytiva) connected to an ÄKTA™ pure chromatography system and eluted with an imidazole gradient (20 mM Tris-HCl, pH 7.8, 500 mM NaCl, 250 mM imidazole, 10% glycerol, and 3.5 mM β-mercaptoethanol).

### Immunoblotting

Purified recombinant ATGL (1.5 µg per lane) was separated by SDS-PAGE and either transferred onto PVDF membranes (0.45 µm pore size; Carl Roth, Karlsruhe, Germany) using wet transfer or analysed by Coomassie Brilliant Blue R250 staining. Membranes were incubated with anti-Xpress® primary antibody (1:5,000; R910-25, Thermo Fisher Scientific), followed by an HRP-conjugated anti-mouse secondary antibody (1:10,000; NA931V, Cytiva). Immunoreactive signals were detected using Clarity Western ECL substrate (Bio-Rad, Hercules, CA, USA) and imaged with a ChemiDoc Imaging System (Bio-Rad). Protein purity was assessed by Coomassie Brilliant Blue R250 staining of a parallel SDS-PAGE gel (Supplementary Fig. S5).

### TAG hydrolase activity assay

ATGL activity toward defined TAG substrates (Larodan, Solna, Sweden; Supplementary Table S1) was determined using purified recombinant ATGL. Substrate emulsions contained 100 µM TAG and 50 µM 1-palmitoyl-2-oleoyl-sn-glycero-3-phosphocholine (Avanti Polar Lipids, Alabaster, AL, USA) in PPB (pH 7.0) supplemented with 2% FA-free BSA (Sigma-Aldrich, St. Louis, MO, USA). Enzymatic reactions were initiated by combining 25 µl substrate emulsion with 2.5 µg purified ATGL in buffer H (0.25 M Sucrose, 1mM DTT, 1mM EDTA), with or without 200 ng purified ABHD5, in a final volume of 50 µl. Reactions were incubated for 90 min at 37 °C. Lipids were extracted by addition of 1 ml methyl-tert-butyl ether/methanol (3:1, v/v) containing 500 nmol butylated hydroxytoluene, 0.1% acetic acid, and 133 pmol internal lipid standard (IST) DAG 34:0-17:0. Following addition of 150 µl water, samples were vortexed and subjected to phase separation by centrifugation (10,000 × g, 10 min). The upper organic phase (700 µl) was collected, dried under nitrogen, and reconstituted in 50 µl 2-propanol/methanol/water (70:25:10, v/v/v). DAG reaction products were quantified by UHPLC-MS/MS using 5 µl of the reconstituted extract.

### UHPLC-MS/MS measurement

DAG species were analysed by UHPLC-MS/MS using an Infinity II/6470 triple quadrupole mass spectrometer (Agilent, Waldbronn, Germany) coupled to an ACQUITY UPLC BEH C18 column (1.7 µm particle size, 2.1 mm inner diameter, 150 mm length; Waters, Milford, MA, USA). The column was maintained at 50 °C. Mobile phases consisted of solvent A (water) and solvent B (isopropanol) containing 0.1% formic acid, 10 mmol l^-1^ ammonium acetate, and 7.7 µmol l^-1^ phosphoric acid. Chromatographic separation was achieved using the following gradient: solvent B was maintained at 50% from 0 to 0.5 min, increased linearly to 80% at 9 min, and to 100% at 22 min, maintained until 24.5 min, and returned to initial conditions at 25.0 min, followed by re-equilibration until 30.0 min. The flow rate was set to 150 µl min^-1^. Analytes were detected in positive ion mode using dynamic multiple reaction monitoring (dMRM). Source parameters were as follows: capillary voltage, 3500 V; gas flow 5 l min^-1^, nebulizer pressure, 30 psi; sheath gas temperature, 400 °C; and sheath gas flow, 12 l min^-1^. MRM transitions, collision energies, and retention time windows for individual analytes are provided in Supplementary Table S2. Analyte peaks were integrated using Skyline (version 26.1.0.057). DAG species were quantified relative to IST DAG 17:0-17:0 using single point calibration and expressed as arbitrary units per IST per mg protein per hour (AU IST^-1^ mg^-1^ protein h^-1^).

## Data availability

The Boltz-2 predictions pipeline, PAE/PDE/pLDDT tensors, mmCIF structural models, *F*_score_ calculation code, and analysis CSV files generated in this study will be made available in a GitHub repository upon publication.

## Acknowledgements

We thank Dr Mariana Colaço-Gaspar Hartig, Dr Peter Hofer, and Dominik Prem, BSc, MSc, for their technical support with protein purification using the ÄKTA™ pure chromatography system. We are also thankful to Dr Robert Zimmermann and Dr Achim Lass for critically reviewing the manuscript.

## Author contributions

A.S.S. conceived the initial study concept and developed the computational approach.

A.S.S and F.P.W.R. jointly developed the overall research strategy, with F.P.W.R. developing the biochemical experimental approach. Both supervised the work. N.B.E. planned and performed all experimental work, including the enzymatic validation assays, and contributed to drafting and editing the manuscript. C.G. and K.G. provided structural biology input and advice on the computational analyses. T.Z. supervised the mass spectrometric measurements and their analysis. A.S.S. implemented the computational methodology.

A.S.S. and F.P.W.R. drafted the manuscript. All authors reviewed and edited the manuscript.

## Materials & Correspondence

The data supporting the findings of this study are presented within the article and its supplementary materials. Additional information or raw data are available from the corresponding authors upon reasonable request.

