## supplementary data for "Triacylglycerol fatty acid composition modulates ATGL regioselectivity and diacylglycerol output"

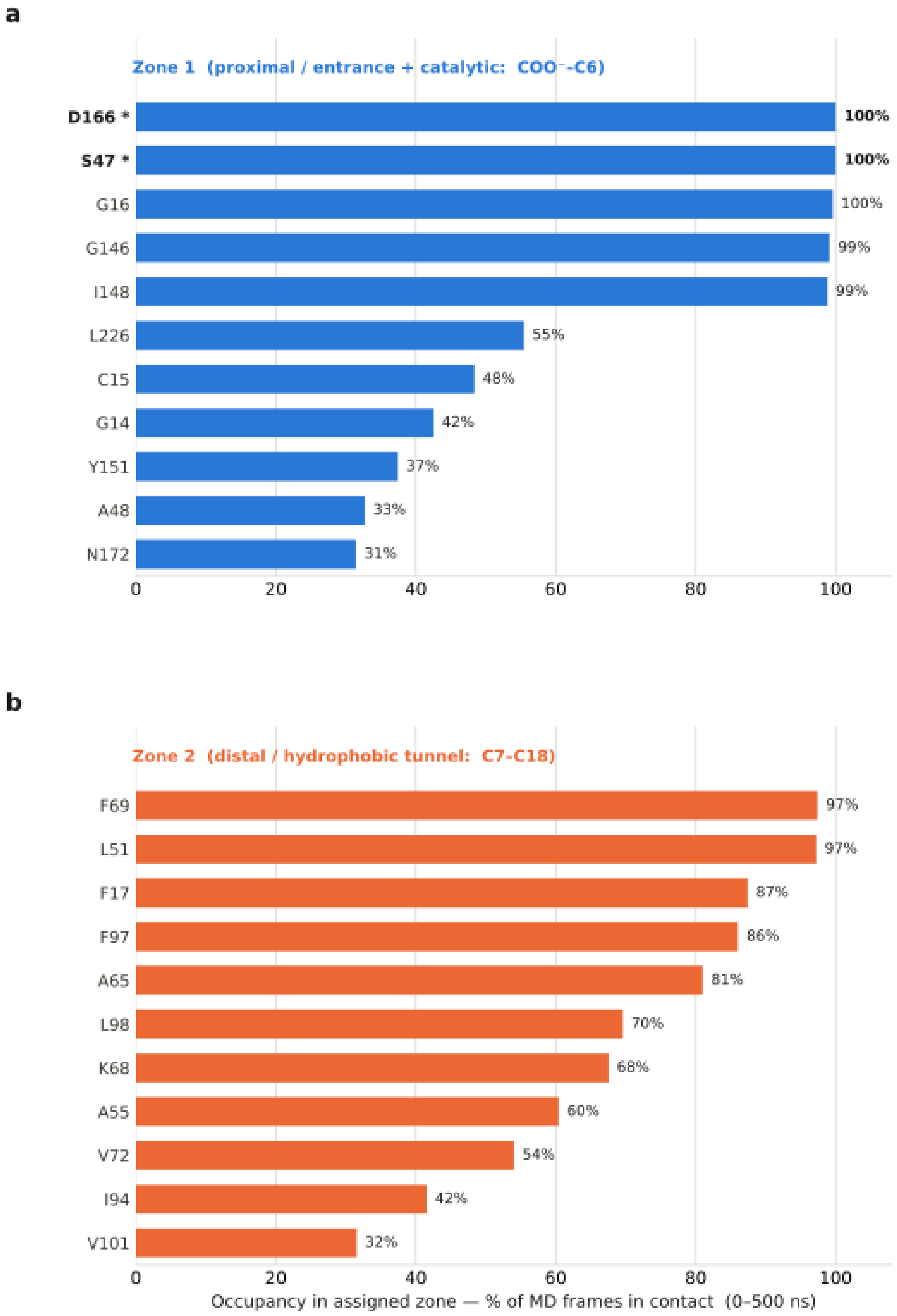


**Figure S1: ATGL–stearic acid contact map over 500 ns of unbiased molecular dynamics.** Contact occupancies were calculated over the full 500 ns production trajectory. A residue–ligand contact was defined as a periodic-boundary-condition (PBC)-aware minimum distance

≤4.0 Å between any residue heavy atom and any ligand heavy atom. The stearic acid ligand was divided into the carboxyl group (C1, O1, and O2) and the acyl chain (C2–C18). Residues with a contact occupancy >30% were assigned to either the hydrophilic anchor (C1−C6) or the hydrophobic acyl tunnel (C7−C18), based on the region in which they showed the greater fraction of contacting frames. Bars show contact occupancy in the assigned region, expressed as the percentage of frames in which the residue contacted at least one ligand atom within that region. (**a**) Residues assigned to the hydrophilic anchor (blue). The catalytic residues S47 and D166 (bold, asterisks) and the oxyanion-hole backbone region (G14–G16) contacted the carboxyl group with occupancies of 99–100%, consistent with a stably substrate-binding configuration. G16 is the only residue with substantial occupancy in both regions (47% in the acyl tunnel) and was assigned to the hydrophilic anchor based on its higher occupancy there (100%). (**b**) Residues assigned to the hydrophobic acyl tunnel (orange). The distal portion of the acyl chain contacted predominantly hydrophobic residues, including F69, L51, F17, F97, A65, L98, V72, I94, V101, and the hydrophobic portion of K68. The contact pattern therefore separates into a hydrophilic anchor around the carboxyl group and a predominantly hydrophobic acyl tunnel surrounding the C7–C18 portion of the acyl chain.

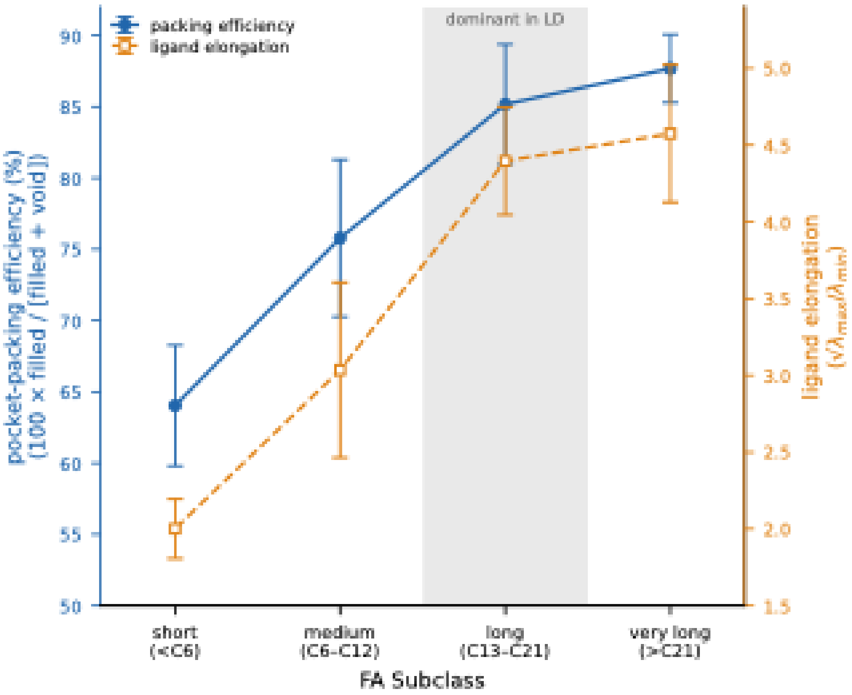


**Figure S2: Binding-pocket packing efficiency and ligand elongation increase with fatty acyl chain length during molecular dynamics (MD).** Pocket-packing efficiency (blue, left axis) and ligand elongation (orange, right axis) are shown as mean ± SD of pooled per-frame values for each fatty acid (FA) subclass. Both metrics were calculated frame-by-frame over 100 ns of production MD for each FA using pyKVFinder with as solvent-excluded surface (SES), grid step of 0.4 Å. probe-in radius of 1.4 Å, probe-out radius of 6.0 Å, and ligand cutoff of 5.0 Å. Packing efficiency was calculated as 100 × filled/(filled + void), where 'filled' corresponds to the ligand SES volume within the cavity and void to the remaining cavity volume. Ligand elongation was calculated as the anisotropy of the ligand SES voxel cloud, $\sqrt{(\lambda max/\lambda min)}$, where λmax and λmin are the largest and smallest eigenvalues of the covariance matrix of the SES voxel coordinates (heavy atoms only; hydrogens excluded). A value of 1 indicates an isotropic shape, with higher values indicating a more elongated shape. The long-chain FA subclass, which constitutes the major fraction of the lipid-droplet triacylglycerol substrate pool, is shaded. Packing efficiency increased from 64% for short-chain FAs to approximately 88% for long- and very-long-chain FAs, whereas ligand elongation increased from 2.0 to 4.6 with increasing chain length.

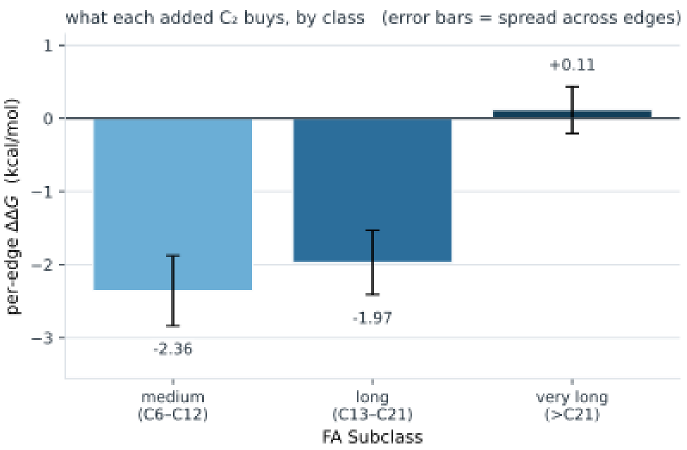


**Figure S3: Relative binding free energy of fatty acid chain elongation across fatty acid subclasses.** Bars show the mean ΔΔG_bind of the C(n)→C(n+2) transformations whose product falls within each subclass: medium-chain C6–C12 (4 edges: C4→C6, C6→C8, C8→C10, and C10→C12), long-chain C13–C21 (4 edges: C12→C14, C14→C16, C16→C18, and C18→C20) and very-long-chain >C21 (2 edges: C20→C22 and C22→C24). No short-chain (<C6) bar is shown because the first transformation in the ladder produces C6. Negative values indicate that chain elongation is energetically favourable. Each edge value is the antisymmetric average, ½(ΔΔG_fwd − ΔΔG_rev), of the forward and reverse TI/MBAR calculations; error bars represent the population standard deviation across edges within each subclass.

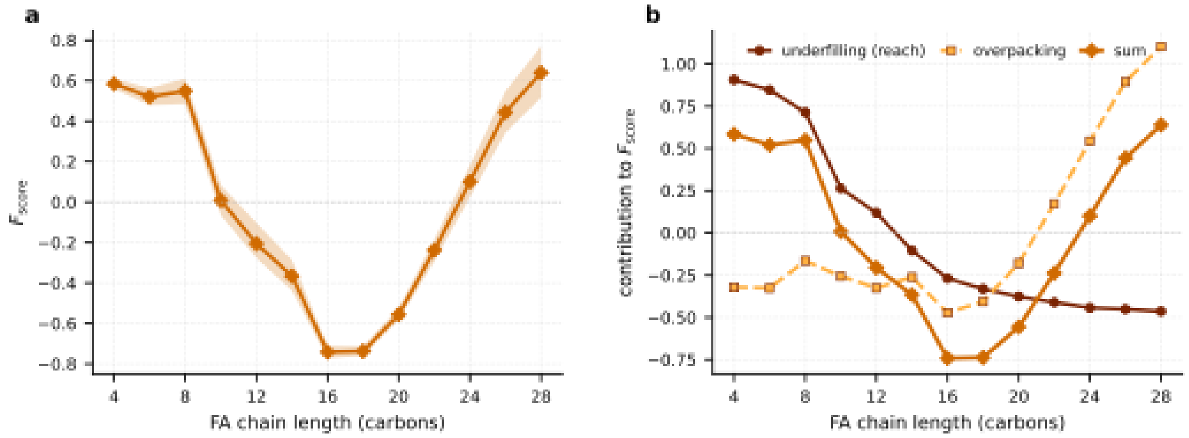


**Figure S4: Chain-length fitness score across the saturated fatty acyl chain series**. Chain-length fitness scores ($F_{score}$) were calculated for the saturated fatty acid (FA) series from C4:0 to C28:0 at even chain lengths. (**a**) $F_{score}$ values with the shaded band indicating the 5th–95th percentile interval from 4,000 bootstrap replicates, in which the 25 Boltz-2 models generated for each FA were resampled with replacement. (**b**) Decomposition of the $F_{score}$ into its two weighted components: underfilling (dark brown, circles, solid line) and overpacking (light amber, squares, dashed line), together with their weighted sum (mid brown, diamonds), which corresponds to the $F_{score}$ shown in (a).

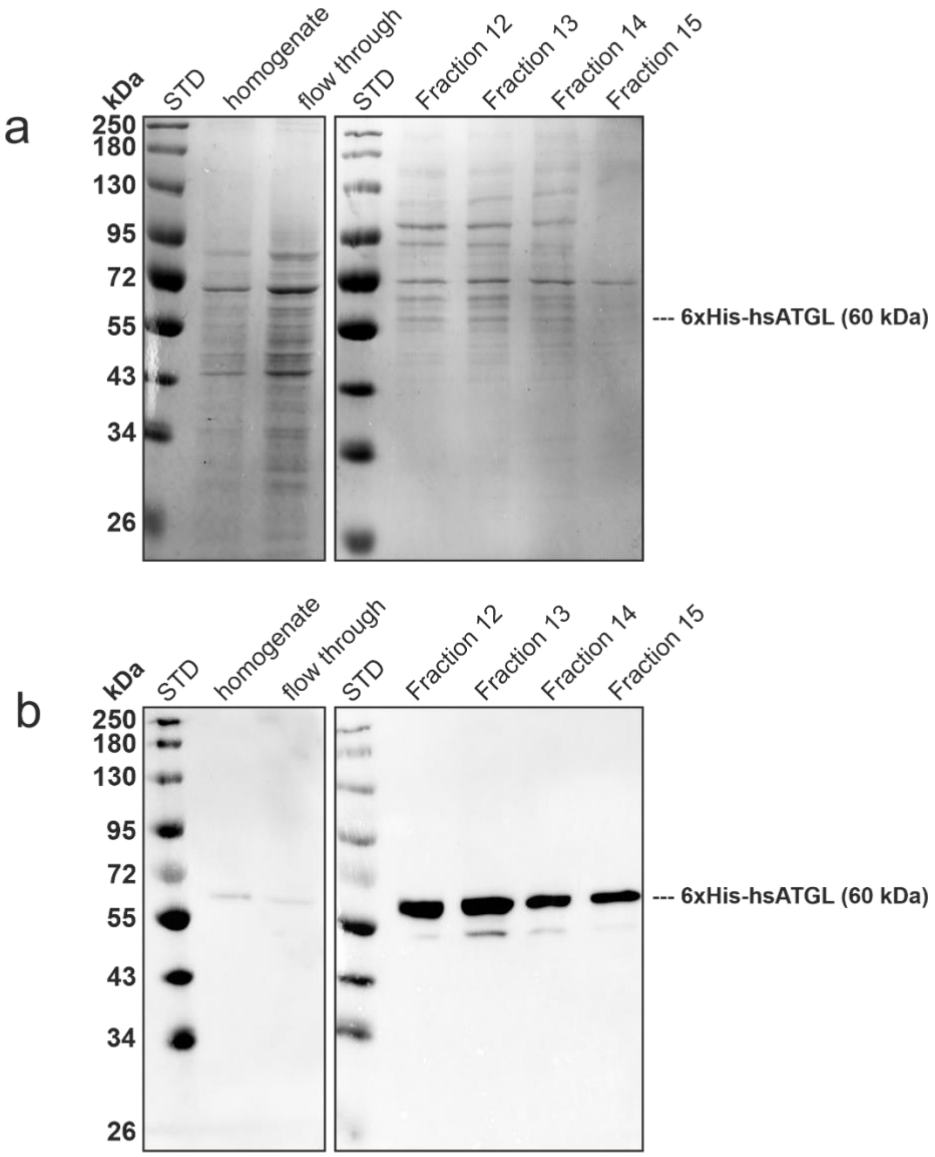


**Figure S5: Purification of recombinant human ATGL.** (**a, b**) Aliquots (1.5 µg) of fractions 12–15 and 1 µl of homogenate and flow-through controls were analysed by 10% SDS–PAGE. “STD” denotes the protein standard (Color Protein Standard, Broad Range; NEB). “Homogenate” denotes the cell lysate before loading onto the HisTrap column, whereas “flow-through” denotes the unbound material collected after column loading. Fractions 12–15 correspond to 2-ml elution fractions collected by the fraction collector. (**a**) Coomassie Brilliant Blue R-250 staining was used to assess protein purity and showed enrichment of His-tagged human ATGL (6×His-hsATGL) in fractions 12–15. (**b**) Following SDS–PAGE, proteins were transferred to a PVDF membrane by wet transfer and 6×His-hsATGL was detected using an anti-Xpress antibody. The recombinant protein was highly enriched in the elution fractions.

**Table 1: TAG substrates used in ATGL activity assays.** All TAG substrates were purchased from Larodan (Solna, Sweden). Catalogue numbers are provided for reference.

| Common name | Fatty acyl composition (*sn*-1/*sn*-2/*sn*-3) | Catalogue no. |
|---|---|---|
| 1,3-Butyrin-2-olein-*sn*-glycerol | 4:0/18:1/4:0 | 34-0418 |
| 1,3-Olein-2-butyrin-*sn*-glycerol | 18:1/4:0/18:1 | 34-1844 |
| 1,3-Laurin-2-olein-*sn*-glycerol | 12:0/18:1/12:0 | 34-1229 |
| 1,3-Olein-2-laurin-*sn*-glycerol | 18:1/12:0/18:1 | 34-1819 |
| 1,3-Palmitin-2-olein-*sn*-glycerol | 16:0/18:1/16:0 | 34-1611 |
| 1,3-Olein-2-palmitin-*sn*-glycerol | 18:1/16:0/18:1 | 34-1831 |
| 1,3-Tetracosanoin-2-olein-*sn*-glycerol | 24:0/18:1/24:0 | 34-2418 |
| 1,3-Olein-2-lignocerin-*sn*-glycerol | 18:1/24:0/18:1 | 34-1823 |
| 1-Stearin-2-olein-3-palmitin-*sn*-glycerol | 18:0/18:1/16:0 | customized |

**Table 2: UHPLC-MS/MS transitions and instrument parameters for the quantification of DAG species.**

| Lipid | Precursor [m/z] | Product [m/z] | RT [min] | RTW [min] | Fragmentor [V] | CAV [V] | CE [V] | Precursor Adduct | Cycle Time [ms] |
|---|---|---|---|---|---|---|---|---|---|
| DAG 8:0-4:0 | 250,2 | 145,1 | 10 | 20 | 110 | 5 | 19 | [M+NH4]1+ | 734 |
| DAG 22:1-18:1 | 444,4 | 145,1 | 15 | 20 | 110 | 5 | 19 | [M+NH4]1+ | 734 |
| DAG 22:1-4:0 | 444,4 | 339,3 | 15 | 20 | 110 | 5 | 19 | [M+NH4]1+ | 734 |
| DAG 24:0-12:0 | 474,4 | 257,2 | 15 | 20 | 110 | 5 | 19 | [M+NH4]1+ | 734 |
| DAG 30:1-12:0 | 556,5 | 339,3 | 15 | 20 | 110 | 5 | 19 | [M+NH4]1+ | 734 |
| DAG 30:1-18:1 | 556,5 | 257,2 | 15 | 20 | 110 | 5 | 19 | [M+NH4]1+ | 734 |
| DAG 32:0-16:0 | 586,5 | 313,3 | 17 | 20 | 110 | 5 | 19 | [M+NH4]1+ | 734 |
| DAG 34:0-16:0 | 614,6 | 341,3 | 17 | 9 | 110 | 5 | 19 | [M+NH4]1+ | 734 |
| DAG 34:0-18:0 | 614,6 | 313,3 | 17 | 9 | 110 | 5 | 19 | [M+NH4]1+ | 734 |
| DAG 34:1-16:0 | 612,6 | 339,3 | 17 | 9 | 110 | 5 | 19 | [M+NH4]1+ | 734 |
| DAG 34:1-18:1 | 612,6 | 313,3 | 17 | 9 | 110 | 5 | 19 | [M+NH4]1+ | 734 |
| DAG 36:1-18:0 | 640,6 | 339 | 17 | 9 | 110 | 5 | 19 | [M+NH4]1+ | 734 |
| DAG 36:1-18:1 | 640,6 | 341,3 | 17 | 9 | 110 | 5 | 19 | [M+NH4]1+ | 734 |
| DAG 36:2-18:1 | 638,6 | 339 | 16,9 | 9 | 110 | 5 | 19 | [M+NH4]1+ | 734 |
| DAG 42:1-18:1 | 724,7 | 425,4 | 20 | 9 | 110 | 5 | 19 | [M+NH4]1+ | 734 |
| DAG 42:1-24:0 | 724,7 | 339,3 | 20 | 9 | 110 | 5 | 19 | [M+NH4]1+ | 734 |
| DAG 48:0-24:0 | 810,8 | 425,4 | 20 | 15 | 110 | 5 | 19 | [M+NH4]1+ | 734 |
| DAG 34:0-17:0 | 614,6 | 327,3 | 16,6 | 9 | 110 | 5 | 19 | [M+NH4]1+ | 734 |

**RT = retention time; RTW = retention time window; CAV = cell accelerator voltage; CE = collision energy**
